# Two spaced training trials induce associative ERK-dependent long-term memory in *Neohelice granulata*

**DOI:** 10.1101/2020.04.16.045427

**Authors:** Santiago Ojea Ramos, Matías Andina, Arturo Romano, Mariana Feld

**Affiliations:** Departamento de Fisiología, Biología Molecular y Celular - Facultad de Ciencias Exactas y Naturales - Universidad de Buenos Aires. Instituto de Fisiología, Biología Molecular y Neurociencias (IFIBYNE) - UBA-CONICET. Ciudad Universitaria (C1428EHA). Buenos Aires, Argentina

**Keywords:** CRAB, SPACING EFFECT, CONSOLIDATION, RECONSOLIDATION, PD 98059, MAPK

## Abstract

Memory formation depends upon several parametric training conditions. Among them, trial number and inter-trial interval (ITI) are key factors to induce long-term retention. However, it is still unclear how individual training trials contribute to mechanisms underlying memory formation and stabilization. Contextual conditioning in *Neohelice granulata* has traditionally elicited associative long-term memory (LTM) after 15 spaced (ITI = 3 min) trials. Here, we show that LTM in crabs can be induced after only two training trials by increasing the ITI to 45 min (2t-LTM) and maintaining the same training duration as in traditional protocols. This new LTM observed was preserved for at least 96 h, exhibited protein synthesis dependence during consolidation and reconsolidation as well as context-specificity. Moreover, we demonstrate that 2t-LTM depends on inter-trial and post-training ERK activation showing a faster phosphorylation after the second trial compared to the first one. In summary, we present a new training protocol in crabs with reduced number of trials that shows associative features similar to traditional spaced training. This novel protocol allows intra-training manipulation and the assessment of individual trial contribution to LTM formation.

## 1. Introduction

Memory processes require the activation of several transduction pathways that lead to post-translational modifications of proteins and gene expression regulation (Dudai, 2012), thus promoting the stabilization of the memory trace. Training protocols that integrate multiple trials spaced over time are more effective in the induction of long-term memory (LTM) than protocols involving massed presentation of training trials. This ubiquitous behavioral occurrence known as spacing effect was first described by Ebbinghaus (1885), and has since been observed in different species (Aziz et al., 2014; Bello-Medina et al., 2013; Pagani et al., 2009; Philips et al., 2007; Toda et al., 2009; Vlach et al., 2008). LTM formation has been shown to be induced by specific time intervals between trials in very different learning and memory animal paradigms including vertebrates (Bolding & Rudy, 2006; Klapdor & van der Staay, 1998; Williams et al., 1991) as well as invertebrates (Carew et al., 1972; Gerber et al., 1998; Lukowiak et al., 1998; Maldonado et al., 1997; Rogers et al., 1994; Tully et al., 1994). However, while the spacing effect has been thoroughly characterized from a behavioral standpoint very little is known about the molecular and synaptic mechanisms that underlie this phenomenon. Work addressing this question in the mollusk *Aplysia californica* showed that there is a narrow permissive window (45 min) on the spacing of two-trial training in order to induce long term sensitization (LTS) (Philips et al., 2007, 2013). This memory process seems to be mediated by extracellular signal-regulated kinase/mitogen-activated protein kinase (ERK/MAPK) phosphorylation and nuclear translocation. Interestingly, activation of ERK by the first trial was not sufficient to induce LTS and the second trial was necessary to prompt it, suggesting that the molecular machinery recruited by the first trial is required to interact with that triggered by the second trial in order to elicit LTS.

In line with this notion, work in *Drosophila melanogaster* proposed that spaced training protocols generate repetitive waves of MAPK activation defined by the duration of inter-trial intervals (ITIs), while massed training induced only one peak of MAPK activation after the last stimulus (Pagani et al., 2009). Accordingly, spaced training induced MAPK-dependent activation of c-Fos and CREB, two known ERK targets involved in LTM consolidation (Cammarota et al., 2000; Dash et al., 1995), but not after massed training (Miyashita et al., 2018). These results support the idea that spaced training is more effective than massed training due to recruitment of specific molecular and cellular mechanisms shown to support LTM (Naqib et al., 2012).

A large body of work has extensively studied the memory processes and their molecular basis on the crab *Neohelice granulata* (formerly *Chasmagnathus granulatus*) (Feld et al., 2005, 2008; Frenkel et al., 2002; Freudenthal & Romano, 2000; Locatelli & Romano, 2005; María Eugenia Pedreira et al., 2004; María Eugenia Pedreira & Maldonado, 2003; Arturo Romano et al., 2006; Tomsic et al., 2003). The behavioral approach to study LTM of the crab *Neohelice* takes advantage of the crab’s innate escape response elicited by the presentation of a visual danger stimulus (unconditioned stimulus, US). Training sessions typically consist of 15 trials of CS-US presentation spaced by 3 min ITI (Fustiñana et al., 2013; Maldonado, 2002) and LTM is robustly evidenced 24 h later by a decrease on the escape response that can last for up to 5 days. LTM is protein synthesis-dependent, context-specific, sensitive to labilization/reconsolidation and extinction. However, the number of trials needed and the relatively short ITI pose a complication when studying the individual trial contribution to molecular mechanisms. Here, we present a newly developed training protocol consisting of only two conditioning trials spaced by 45 min that elicits robust LTM and offers a valuable tool for studying the single contribution of individual trial to memory formation. We further show this memory to be context-specific, protein synthesis-dependent and mediated by ERK activation. Memory reactivation upon a unique CS presentation 24 h after training session suggest two-trial elicited LTM (2t-LTM) can be rendered labile again and must undergo reconsolidation to restabilize while protein synthesis inhibition blocked memory expression 24 h afterwards. Finally, we assessed 2t-LTM ERK kinetics and discussed the molecular implications of the spacing effect observed in different tasks, including learning and memory in *Neohelice*. Altogether, our results show that 2t-LTM improves the possibility of pharmacological manipulations during the 45 min ITI and provides insight on individual trial input to unravel the molecular mechanisms behind the spacing effect.

## 2. Material and Methods

### 2.1. Animals

Adult male *Neohelice granulata* measuring 2.7–3.0 cm across the carapace and weighing an average of 14.97 ± 0.47g were collected from narrow coastal inlets of San Clemente del Tuyú, Argentina, and transported to the laboratory where they were housed in plastic tanks (32 × 46 × 20 cm, 20 animals per tank) filled to a depth of 1 cm with 12 ‰ diluted seawater (prepared from Cristalsea Marinemix salts, USA).

The holding and experimental rooms were kept on a 12 h light-dark cycle (lights on from 08:00 am to 20:00 pm) and the temperature was set on a range of 22-24 °C. Experiments were carried out within the first week after capture and each crab was used in only one experiment. Animals were maintained and experiments were conducted in accordance with the National Research Council Guide for the Care and Use of Laboratory Animals (USA) as well as Argentinean guidelines for ethical use of laboratory animals.

### 2.2. Drugs and injection procedure

Dimethyl sulfoxide (DMSO, Anedra, Argentina) was used as vehicle (VEH). The VEH or drug solutions were injected through the right side of the dorsal cephalotoracic-abdominal membrane using a Hamilton syringe with a needle fitted with a plastic cannula to control the penetration depth to 4 mm, ensuring that the desired solution is injected into the pericardial sac. The total volume of haemolymph in a crab has been estimated in 5 ml (30 % of the body weight; Gleeson and Zubkoff, 1977) and 10ul of VEH or drug were injected, resulting in a dilution of approximately 500-fold for DMSO of injected drugs.

The following drugs were used:

A stock solution (74.8 mM) of the MEK inhibitor 2-(2-amino-3-methoxyphenyl)-4H-1-benzopyran-4-one (PD98059, PD hereafter, Sigma-Aldrich, Argentina) was conserved at −20 °C and freshly diluted in DMSO to the desired concentration (11.22 mM, final dose of 1.765 μg/g) the day of the experiment. The protein synthesis inhibitor Cycloheximide (CHX, Sigma-Aldrich, Argentina) was diluted in DMSO on the day of the experiment and 10 μl were injected (40 μg/crab, a dose commonly used at our lab, Pérez-Cuesta & Maldonado, 2009).

### 2.3. Experimental device

The experimental device used to train and to test animals, called actometer, consists of a bowl-shaped plastic container where one crab is placed. The unconditioned stimulus (US) consists of an opaque rectangular screen that moves horizontally over the animal and provokes the crab’s innate running response (Maldonado et al., 1997). Two light sources allow changing the context by illuminating the actometer from above (upper light) or below (lower light) the container. Vibrations produced by the animal movement were registered by four microphones attached to the base of the container. This signal was amplified, integrated during the entire trial and translated into arbitrary numerical units ranging from 0 to 22,500 by a computer.

### 2.4. Behavioral procedures

Each crab was placed in a container and initially adapted to it during 13 min with lower light. Each trial started with an illumination switch from lower light to upper light (CS), 18 sec prior to the screen movement. Two successive events of horizontal cyclical screen movement drawing a 90 ° angle and lasting 9 sec defined the US ending together with a new light switch from upper to lower. During intertrial intervals (ITIs), the actometers remained illuminated with the lower light. Trial sequences, illumination, duration and ITI were programmed and controlled by the registering computer. The experimental room contained forty experimental devices separated from each other by partitions and allowed training or testing of 40 crabs simultaneously.

Training: Each training session consisted of two trials and a 45 min ITI, unless stated otherwise. In all experiments one group was trained (TR) while the other group underwent the same manipulation but didn’t receive any US (control group, CT). In drug administration experiments two pairs of CT and TR groups were injected with either drug or VEH. Animals were randomly assigned to the experimental groups.

Reactivation: Twenty-four hours after training animals were placed in the experimental device and after the adaptation period the upper lights turned on for 27 sec, without US presentation. This protocol has been found effective to trigger labilization of the consolidated LTM (Fustiñana et al., 2013).

Testing: Testing for LTM began 24 h after training or reactivation. During the testing session animals were placed in the actometer for 10 min adaptation and later received three spaced trials (ITI: 153 sec).

### 2.5. Procedure for ERK/MAPK phosphorylation determination

Animals were anaesthetized by immersion in iced marine water for five minutes and central brain dissection was performed. Each central brain was homogenized with 15 strokes in a Dounce Tight homogenizer with 50 μl of Buffer A (10 mM Hepes pH 7.9; 1,5 mM MgCl2; 10 mM KCl; 1 mM DTT; 1 μg/ml Pepstatin A; 10 μg/ml Leupeptin; 0,5 mM PMSF; 10 μg/ml Aprotinin; 1 mM sodium orthovanadate and 50 mM sodium fluoride). The homogenate was centrifuged for 15 min at 16000 x *g* and the supernatant was aliquoted and kept at −20 °C until used. All the extraction protocol was performed at 4 °C.

Fifteen μl of sample were run in 12.5% SDS-PAGE gels at 100V for 2 h and then blotted to 0.45 μm low-fluorescence PVDF membrane (Millipore-Merck, USA) at 100V for 50 min. Membranes were then blocked in 5% nonfat dry milk in Tris Buffered Saline-0.1 % Tween 20 (TTBS) and incubated with primary antibodies against phospho-ERK (pERK, Santa Cruz Biotechnology, sc-7383, 1:500) and total ERK (ERK, Cell Signaling Technologies, cat. #9102, 1:1000) in TTBS at 4 °C. Detection was performed using IRDye secondary antibodies (Li-Cor, USA) at a 1:10,000 dilution, using LI-COR Odyssey Imaging System. Secondary antibodies used were 800CW Donkey anti-Mouse IgG (cat. 926-32212) and 680RD Donkey anti-Rabbit IgG (cat. 926-68073).

### 2.6. Data Analysis

LTM retention criterion was established as a statistically lower normalized response of the trained group compared against control groups in the first trial of the testing session. The escape response during each trial was normalized within animal, against each animal’s maximum escape response. Therefore, the escape response during testing varied between the values 0 and 1. This procedure was done to account for differences between animals due to factors other than training (e.g. weight, stamina, overall health).

Since behavioral data violated normality and heteroscedasticity assumptions Mann-Whitney U test or Kruskal-Wallis test were used followed by Dunn’s Multiple Comparison post-hoc tests. Unless stated otherwise, median +/− interquartile ranges of normalized escape response is shown.In ERK/MAPK phosphorylation assays, Relative Optical Density (ROD) was quantified using NIH ImageJ v1.51j8 software (Schneider et al., 2012). Statistical analysis of the data consisted of ANOVA followed by Holm-Sidak’s multiple *post hoc* comparisons between TR and CT groups.

All data analysis was performed using GraphPad Prism 8 software.

## 3. Results

### 3.1. Two training trials spaced by 45 min elicit LTM formation

The traditional training protocol of contextual pavlovian conditioning (CPC) in *Neohelice granulata* involves 15 trials spaced by 3 min (Fustiñana et al., 2013). Compelling evidence supports the notion that given the adequate ITI, training protocols involving fewer trials would be as effective as higher numbers of stimuli to elicit LTM formation (Philips et al., 2007). Thus, we addressed the question whether two trials were enough to induce LTM in *Neohelice granulata.* Given the drastic reduction in the number of training trials, we examined whether increasing the ITI would lead to LTM formation, as described in previous studies. Therefore, we trained groups of animals with two trials (CS-US presentation) using three different ITIs (3, 45 and 60 min). We also trained a CPC group with 15 trials and 3 min ITI that served as a control for LTM (Fig. 1A). All trained groups (TR groups) were run in parallel to a control group (CT groups), subjected to the same experimental conditions but without US presentation. All groups were tested 24 h after training for LTM.

**Figure 1:**
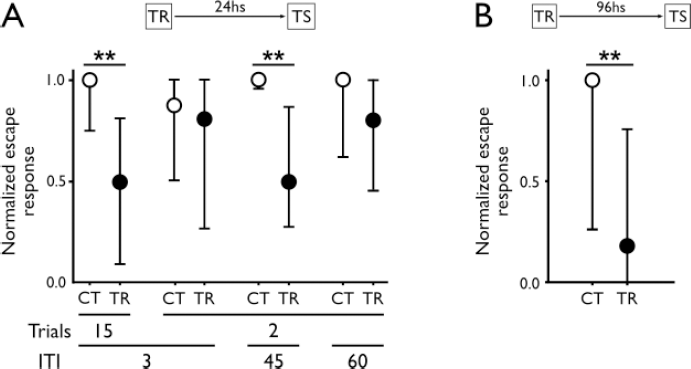
Two trials spaced by 45 min induce LTM lasting for at least 96 h. **A)** Normalized escape response in testing session (24 h after TR) from animals stimulated (TR groups, filled circles) or not (CT groups, empty circles) with the US using four different protocols: 15 trials with 3 min ITI or 2 trials with different ITIs (3; 45 or 60 min). Kruskal-Wallis (H = 23.38, df = 7, p = 0.0015, n = 16 - 20 animals per group) followed by post hoc Dunn’s multiple comparisons test. **p<0.01. **B)** Normalized escape response in testing session (96 h after TR) from animals trained with 2 trials and 45 min ITI. Mann-Whitney test (U = 68, **p<0.01; n=16 - 18 animals per group).

In line with earlier work from our lab (Fustiñana et al., 2013), CPC-trained animals (15 trials, 3 min ITI) showed memory retention during the testing session, evidenced by the significant lower escape response observed between this group and its respective control (Fig. 1A; CT vs TR, p = 0.0016). Strikingly, two-trial training were sufficient to elicit LTM when the ITI was 45 min (CT vs TR, p = 0.0110; from now on referred to as 2t-LTM) but not when they were spaced by 3 (CT vs TR, p > 0.9999) or 60 min (CT vs TR, p = 0.9770), consistent with previous findings on spacing effect.

To address the stability of the 2t-LTM, we tested memory retention 96 h after training. Interestingly, LTM was preserved 96 h after training (Fig. 1B, p = 0.0061), suggesting 2t-LTM strength is conserved in spite of the reduction in trial number.

### 3.2. 2t-LTM consolidation depends on protein synthesis

It is well established that LTM consolidation depends on protein synthesis in invertebrates as well as in vertebrates (Alberini, 2008; Pedreira et al., 1996). Thus, we tested whether protein synthesis is also required for 2t-LTM consolidation. Pairs of TR and CT groups were systemically administered with cycloheximide (CHX, a eukaryotic protein synthesis inhibitor) immediately before (Fig. 2A) or after (Fig. 2B) training. Both experiments included a pair of CT and TR groups injected with vehicle (VEH) at the same time points. While both pairs of VEH.TR groups showed robust memory retention (p = 0.0051 and p = 0.0027 for CT vs TR post hoc comparisons from pre- or post-TR injection, respectively), CHX administration impaired 2t-LTM formation when injected both before (p = 0.239 for CT vs TR post hoc test) or after (p = 0.214 for CT vs TR post hoc test) the training session (Fig. 2). These results indicate that, comparable with training protocols of 15 trials with 3 min ITI, 2t-LTM is protein synthesis dependent.

**Figure 2:**
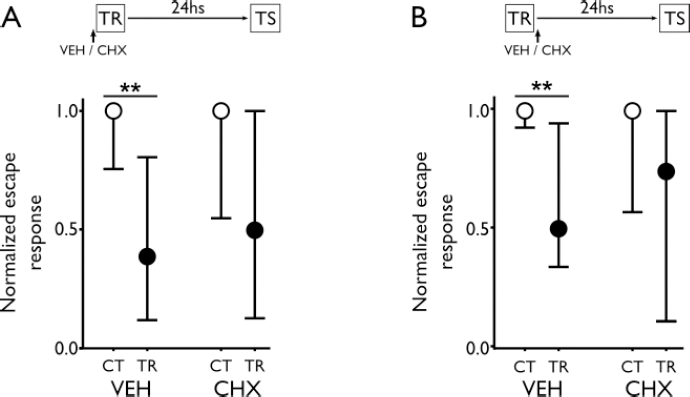
2t-LTM depends on protein synthesis. CHX or VEH were administered immediately pre-TR (H = 13.28, df = 3, p = 0.0041, n = 18 – 20 animals per group, **A**) or post-TR (H = 14.47, df = 3, p = 0.0023, n = 17 – 18 animals per group, **B**) and animals were tested 24hs after TR. Kruskal-Wallis followed by post hoc Dunn’s multiple comparisons test. TR groups, filled circles; CT groups, empty circles; **p<0.01.

### 3.3. Retrieval of 2t-LTM induces the labilization/reconsolidation process

Associative LTMs are capable of being reactivated and labilized upon retrieval (Alberini & Ledoux, 2013). Labile memories are susceptible to disruption or updating (Forcato et al., 2011; Krawczyk et al., 2015; Fustiñana et al., 2014). If no disrupting process follows labilization, the memory becomes restabilized through a process called reconsolidation (Nader et al., 2000; Przybyslawski & Sara, 1997). Moreover, LTM reconsolidation has been demonstrated to be protein synthesis dependent in many animal species (Cai et al., 2012; Fustiñana et al., 2013; Nader et al., 2000; María Eugenia Pedreira et al., 2002).

In order to assess whether 2t-LTM is labilized after retrieval, we inhibited protein synthesis after memory reactivation. Twenty four hours after the training session, two pairs of TR and CT groups of animals were subjected to memory reactivation 20 min after injection of VEH or CHX. A third pair of TR and CT groups was injected and kept in the holding containers, to control for protein synthesis inhibition without reactivation (see schematic experimental design in Fig. 3). Twenty four hours after retrieval session, all groups were tested for LTM retention (Fig. 3). TR animals injected with VEH before re-exposure to the same context (SC.VEH) showed LTM retention when compared with the respective CT group (CT vs TR; p= 0.006; Fig. 3, SC.VEH), indicating that VEH injection did not affect memory reconsolidation. In contrast, CHX-injected animals re-exposed to the training context did not show significant differences in normalized escape response during testing session (CT vs TR; p = 0.187; Fig. 3, SC.CHX). Furthermore, CHX administration without context re-exposure did not affect performance (CT vs TR; p = 0.0021; Fig. 3, NR.CHX), suggesting that 2t-LTM reconsolidation process can be triggered specifically after CS-induced memory reactivation.

**Figure 3:**
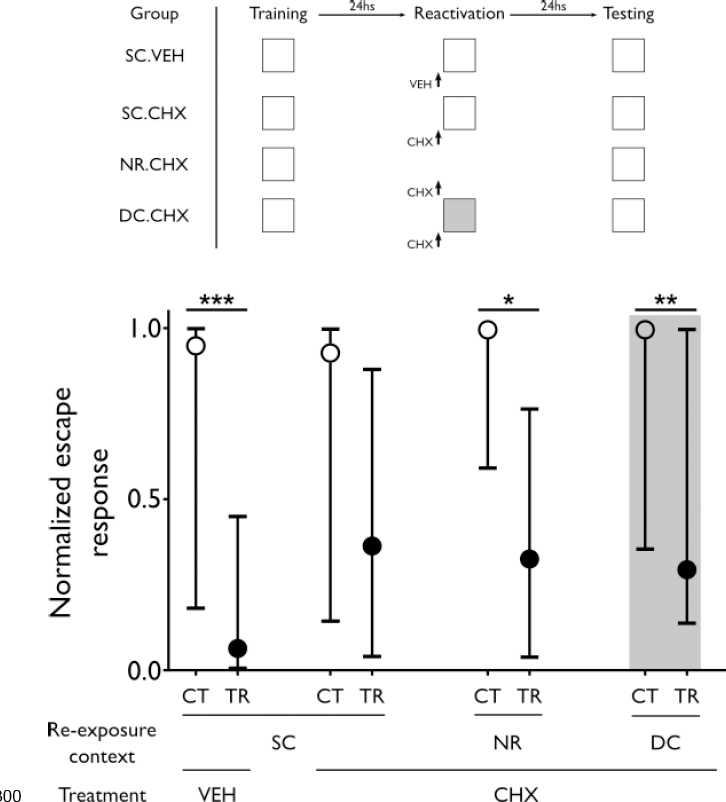
2t-LTM is context-specific and its reconsolidation depends on protein synthesis. CHX or VEH was administered 20 min before memory reactivation in the same context (SC). Pairs of CT and TR animal groups were injected with CHX and 20 min afterwards either subjected to the same manipulation but in a different context (DC) or not re-exposed to any context at all (NR). Schematic experimental design is shown in upper panel. Kruskal-Wallis (H = 42.88, df = 7, p <0.0001, n = 33 - 40 animals per group) followed by post hoc Dunn’s multiple comparisons test. TR groups filled circles; CT groups, empty circles; *p<0.05; **p<0.01; ***p<0.001.

### 3.4. 2t-LTM is context-specific

Several studies have shown that contextual memories can undergo labilization/reconsolidation only when subjects are briefly re-exposed to the training context, while exposure to a different context fails to reactivate the associative memory (de la Fuente et al., 2011; Fustiñana et al., 2013; María Eugenia Pedreira et al., 2002; Suzuki et al., 2004). Given that 2t-LTM can be labilized by re-exposing animals to the training context (Fig. 3, SC.CHX groups), this opens the question whether reactivation of this memory is specific to the TR context where the original CS-US association was established. To address this question, we included a fourth pair of CT and TR groups in the previous experiment which were re-exposed to a different context (DC) 20 min after CHX administration (Fig. 3, DC.CHX groups). We predicted that if 2t-LTM entails a specific association between the context and the US, exposure of animals to a different context 24 h after TR would not labilize the original memory, and CHX injection would not have any effect on memory reconsolidation.

Consistent with our hypothesis that 2t-LTM reactivation is context-specific, memory was not impaired in DC.CHX pair of groups. Normalized escape response from TR group was significantly lower than the corresponding CT group when tested 24 h after being placed in a different context under the effect of the protein synthesis inhibitor (p = 0.043; Fig 3, CHX.DC). Together these results show that 2t-LTM can undergo labilization/reconsolidation only after re-exposure to the same TR context while placement in a different CS (different context) failed to trigger these processes, further supporting context-specificity. Moreover, it discards possible spurious interactions between CS exposure and drug administration.

### 3.5. 2t-LTM depends on ERK/MAPK pathway

LTM in invertebrates has previously been linked to the activation of the MAPK pathway (Crow et al., 1998; Purcell et al., 2003; Sharma et al., 2003). Specifically, ERK involvement in LTM formation has already been described in *Neohelice* (Feld et al., 2005). According to previous reports, 2-trial/45 min-ITI memory in *Aplysia* (Philips et al., 2013) depends on a specific ERK activation kinetics, Consequently, we aimed at dissecting this signaling pathway participation in 2t-LTM.

In order to asses ERK contribution to 2t-LTM formation, we administered PD 98059 (PD), a MEK (ERK kinase) inhibitor, at different times before, during and after training (Fig. 4) at concentrations that had previously been established to efficiently inhibit ERK activity and to block LTM consolidation in *Neohelice granulata* (Feld et al., 2005).

**Figure 4:**
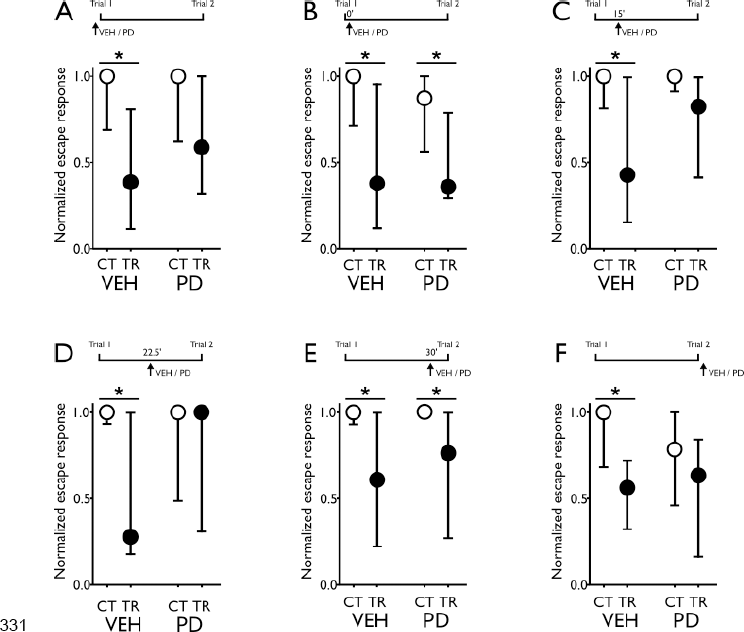
2t-LTM depends on ERK/MAPK activation at specific timepoints. PD 98059 (PD, a MEK inhibitor) or VEH were administered immediately before (H = 14.4, df = 3, p = 0.0024; **A**) or after (H = 10.29, df = 3, p = 0.0163; **F**) training; or immediately after (H = 16.69, df = 3, p = 0.0008; **B**); 15 min (H = 11.93, df = 3, p = 0.0076; **C**); 22.5 min (H = 8.12, df = 3, p = 0.039; **D**) or 30 min (H = 16.38, df = 3, p = 0.0009; **E**) after the first training trial and 24 h later animals were tested for memory retention. Kruskal-Wallis followed by post hoc Dunn’s multiple comparisons test. TR groups, filled circles; CT groups, empty circles; *p < 0.05; **p < 0.01; n = 14-20 animals per group.

Administration of PD both immediately before (Fig. 4A; CT.VEH vs TR.VEH, p = 0.011; CT.PD vs TR.PD, p = 0.098) and after training (Fig. 4F; CT.VEH vs TR.VEH p = 0.032; CT.PD vs TR.PD p = 0.32) had an amnesic effect, supporting ERK requirement during 2t-LTM formation.

However, PD administration at different times during ITI showed distinct effects on memory retention. It impaired LTM when administered at 15 (Fig. 4C; CT.VEH vs TR.VEH, p = 0.010; CT.PD vs TR.PD, p = 0.355) or 22.5 min (Fig. 4D; CT.VEH vs TR.VEH, p = 0.010; CT.PD vs TR.PD, p > 0.999), but not when injected immediately (Fig. 4B; CT.VEH vs TR.VEH, p = 0.007; CT.PD vs TR.PD, p = 0.02) or 30 min (Fig. 4E; CT.VEH vs TR.VEH, p = 0.02; CT.PD vs TR.PD, p = 0.007) after the first trial. Thus, these results are in line with the hypothesis of tight temporal regulation of ERK activation during 2t-LTM formation.

### 3.6. ERK/MAPK pathway activation dynamics in 2t-LTM

In order to understand ERK dynamics in 2t-LTM formation, we assessed ERK activation in protein extracts from the central brain of *Neohelice* obtained at different times during and after training. We measured the phosphorylation levels of ERK by immunoblotting against phosphorylated ERK (pERK) and total ERK independently of its phosphorylation state (tERK). Then, pERK/tERK ratios were calculated and normalized against *naïve* (NV) activation levels.

ERK was significantly activated 30 min after the first trial (Fig. 5A, CT vs TR, p = 0.0004). ERK phosphorylation from TR groups at other time points during ITI were not significantly different from CT or NV groups, suggesting a small outflow from basal levels. CT groups did not differ significantly from NV groups. These results, together with the memory impairment observed by intra-TR PD administration (Fig. 4C and D), support a transient ERK activation window necessary for memory formation after the first trial.

**Figure 5:**
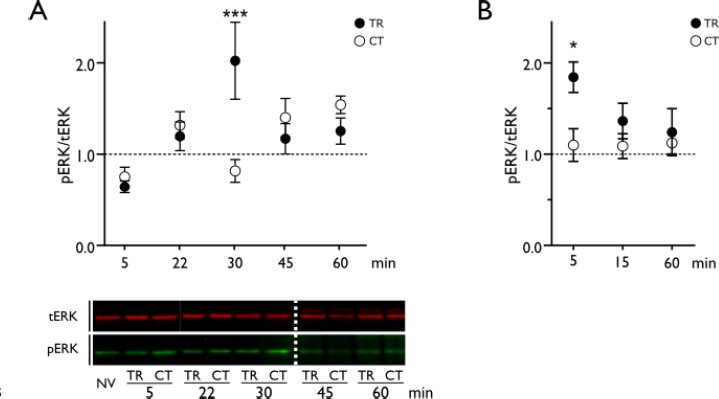
2t-LTM formation ERK/MAPK activation kinetics. Animals were euthanized at different time points after first TR trial (F_10, 86_ = 4.46, p < 0.0001; **A**) or after TR completion (F_6, 57_ = 2.30, p = 0.046, **B**). Mean +/− SEM of Naïve-relativized ERK activation calculated as pERK / totalERK optical density from control (CT, empty circles); trained (TR, filled circles) and naïve (NV, dotted line) animals are shown. Representative tERK and pERK blots are shown (white dotted line separates different membranes). One-way ANOVA followed by Holm-Sidak’s post hoc comparisons test CT vs TR. *p<0.05; ***p<0.001; n = 7-10 animals per group.

Surprisingly, a second training trial induced significant ERK activation only at 5 min after training completion (Fig. 5B, CT vs TR, p = 0.013), while later time points were not significantly different from CT or NV groups (p > 0.05). This result supports the requirement of this kinase activation for consolidation of 2t-LTM in *Neohelice*.

In summary, our data suggest that after the first trial, ERK activity is primed for a second trial to induce a faster dynamic, necessary for 2t-LTM formation. Our novel LTM-inducing training protocol entails a useful tool for intra-trial interventions and enables further insight into the molecular mechanisms that regulates the spacing effect and memory formation.

## 4. Discussion

Here we showed that *Neohelice* associative aversive memory can be unveiled by a training protocol involving 2 trials spaced by 45 min, achieving memory retention for at least 96 h (2t-LTM).

In *Neohelice*, as in many different species, the temporal distribution of training trials drastically affects memory retention (Pereyra et al., 2000). It has been shown that 15 or more spaced trials can induce LTM. Moreover, previous studies in this invertebrate showed a requirement of at least 10 spaced trials to induce 24-h retention. Furthermore, experiments exploring the effect of reducing the duration of the ITI indicated that massed training (e.g. ITI shorter than 27 sec) did not allow associative LTM expression even when the number of trials was significantly increased (e.g. 120; 300 or 1000) as to allow a total stimulation time that equals spaced training duration (Maldonado, 2002). Additionally, it was later shown that 4 or 5 trials with ITI of 3 min did not induce memory retention either (Fustiñana et al., 2013; Romano et al., 1996). Consistently with previous results, 2 trials with an ITI of 3 min did not induce LTM (Fig. 1). However, a small number of trials had never been combined with ITIs longer than 3 min, leading to the idea of a boundary condition for LTM formation (Maldonado, 2002) in this model. Results presented here support the notion that fewer trials are able to induce long-term retention (up to 96 h) as long as the adequate ITI (e.g. 45 min) is applied, suggesting this event to be constrained by or contingent upon the presence of a specific molecular environment. Such protocol would allow unraveling the mechanisms associated with the LTM formation during the training.

Protein synthesis dependency has been shown for different memory paradigms in diverse animal species, and for a variety of learning tasks (Davis & Squire, 1984). 2t-LTM shares parametric features with other well established LTM protocols: it lasts for at least 96 h; it requires protein synthesis to be consolidated; retrieval of consolidated memory after context exposure renders memory labile again and, finally, a change in context during retrieval precludes memory labilization. These findings support an associative nature of the memory trace induced by this new training protocol. Moreover, the ITI used provides a temporal window to pharmacologically dissect the contribution of individual trials to LTM formation and study the spacing effect. Current attempts to explain the spacing effect hypothesize there is a refractory period between trials, such that additional stimuli are ineffective for learning improvement (Smolen et al., 2016). One of these hypotheses proposes that time is necessary for the neural circuitry and molecular machinery to adequately record an event. Thus, time between stimuli would allow the system to regain its ability to respond. Another hypothesis argues that the molecular processes occurring towards the end of training allow the formation of a LTM. Thus, the first trials of a spaced training would have a ‘priming’ effect on synapses, so that subsequent stimulation act to reinforce learning. These hypotheses are not mutually exclusive and further molecular studies will lead to better distinguish between these possibilities. Here, we described a new associative memory protocol, using the same trial structure as in CPC, but reducing to a minimum the number of trials while maintaining the same total training duration. Thus, in *Neohelice* CPC 15 trials are delivered in a 45 min training session, while in 2t-LTM training only 2 trials are applied during the same time lapse, leaving a period of almost 45 min in between with no stimulation. Although these memories appear indistinguishable, our results are not conclusive about what is the contribution of trials between 1 and 15 or whether these are functionally different memories. Ongoing experiments will help clarify this interrogation.

ERK activation has already been shown to be necessary for different phases of memory (Bekinschtein et al., 2008; Besnard et al., 2014; Krawczyk et al., 2015, 2016; Merlo et al., 2018) in different species (Alonso et al., 2002; Atkins et al., 1998; Blum et al., 1999; Crow et al., 1998; Feld et al., 2005; Purcell et al., 2003; Sharma et al., 2003). Among invertebrates, ERK phosphorylation has been linked to LTM in invertebrates such as *Aplysia* (Philips et al., 2007, 2013), *Drosophila* (Miyashita et al., 2018) and *Hermissenda* (Crow et al., 1998). ERK activation in crab’s nervous system had also been demonstrated to be required for LTM formation in *Neohelice* (Feld et al., 2005) using a similar protocol to CPC, but without light changes. In these conditions, ERK is activated in the crab`s central brain 1 h after the training session, while massed training or no stimulation control elicit immediate ERK activation. Inhibition of the pathway 45 min (but not immediately or 1 h) after training impairs performance 24 h later, but not at 4 h, supporting a specific requirement of the pathway in memory consolidation. However, intra-training activation or effect of inhibition on memory performance was difficult to assess using 3-min ITIs.

Using a two-trial training paradigm in *A. californica*, Philips et al. (Philips et al., 2007, 2013) dissected the role of each trial. In *Aplysia* LTS pERK peaks at 45 min after the first trial, just at the same time the second trial is applied. The model posits that the first trial would start the activation of cascades that establish a narrow time window during which a favorable molecular environment allows LTM formation to be triggered by a second trial (Philips et al., 2013). The data presented here is consistent with this proposal. However, during 2t-LTM training, ERK phosphorylation increases significantly by 30 min after the first trial and 15 min before second trial is applied. Supporting its requirement for 2t-LTM formation, pharmacological inhibition using PD effectively impaired behavioral performance only when the drug was injected 15 or 22.5 min (but not 30 min) after the first trial, a few minutes previous to ERK peak activity. Furthermore, PD injection after training also impaired memory retention, suggesting ERK activation 5 min after the second trial is required for LTM formation. Noteworthy, ERK activity showed a specific temporal pattern elicited during and after training. This kinetics profile argues in favor of the importance of the second trial in order to induce LTM formation, while providing support for using longer ITI, as ERK activation 30 min after the first trial enables a second trial to trigger LTM. It is still uncertain what other mechanisms are triggered after ERK activation at this time point. We cannot discard the fact that we are using total protein extraction to determine ERK activation, while activity might be localized in a particular subcellular compartment and consequently is being underestimated (Salles et al., 2015). Consistent with this possibility, the first trial memory trace might be clustered in a restrained group of neurons (Kastellakis et al., 2015), which would make differences difficult to detect with these techniques. Experiments are in progress to assess the role of ERK downstream targets in 2t-LTM. But the strong matching between ERK kinetics and pharmacological alteration of behavior strengthens the idea that ERK would be a key factor in 2t-LTM consolidation and/or acquisition.

ERK activation has been shown to be necessary not only for synaptic plasticity, but also for several forms of learning and memory (Thomas & Huganir, 2004). Studies on dendritic spine formation in cultured hippocampal neurons showed that only repeated depolarization induced sustained activation of ERK that correlated with and was essential for spine formation (Wu et al., 2001). Moreover, multiple cytosolic ERK targets have also been found to be relevant for different plastic processes (Ahn, 2009; Earnest et al., 1996; Gong & Tang, 2006; Kelleher et al., 2004; Kneussel & Wagner, 2013; Schrader et al., 2006), supporting a role for this cascade throughout the neuron. At the molecular level, the increase in effectiveness that is observed behaviorally after spaced training can be translated as a greater effectiveness in recruiting the molecular pathways relevant for memory formation (Naqib et al., 2012). In this sense, it has been found that spaced training is more effective in the recruitment of CREB activation than massed training (Genoux et al., 2002; Josselyn et al., 2001). This protein is essential in the regulation of gene expression and has a key role in plasticity and memory formation processes (Kandel, 2012; Kida & Serita, 2014) and its function in specific areas from mice brain has been shown to influence the probability that individual neurons are recruited into a memory trace (Han et al., 2007). Furthermore, using aversive olfactory association in flies, Miyashita and coworkers recently demonstrated that LTM requires repeated training trials with rest intervals between them in order to induce pERK-dependent transcriptional cycling between c-Fos and CREB (Miyashita et al., 2018). In their report, they also show that training trials suppress ERK activity by recruiting ERK phosphatases (calcineurin and protein phosphatase 1), consistent with the previous proposal that length interval is determined by the activity of SHP2 (protein tyrosine phosphatase 2), associated with increased activity of ERK/MAPK (Pagani et al., 2009). Our results slightly differ from the ones described in *Drosophila* and *Aplysia*, as ERK activity showed a significant increase between trials, suggesting different phosphatase kinetics. However, the nature and kinetics of the hypothetical phosphatase activity remain to be elucidated.

This body of results suggests that the spacing effect can be unraveled through assessment of LTM. Contrary to what we had expected, only two trials (instead of 15) and a longer ITI (45min instead of 3min) maintaining the same training duration induced robust LTM formation and allowed intra-training manipulations without impairing behavioral performance. As shown in different experiments, vehicle administration during ITI did not alter memory expression or impair performance during tests. Moreover, this training protocol allows assessing individual trial contribution to LTM formation processes. This represents a great advantage compared to previous protocols involving a larger number of trials and a shorter ITI. Simultaneously, it contributes to the understanding of relevant trial structure for LTM induction and disentangling the mechanisms associated with LTM formation.

## Abbreviations

2t-LTM: two trial-long term memory
CHX: Cycloheximide
CPC: contextual pavlovian conditioning
CS: conditioned stimulus
CT: control group
ERK: extracellular signal-regulated kinase
ITI: inter-trial interval
LTM: long term memory
LTS: long-term sensitization
MAPK: mitogen-activated protein kinase
PD: PD 98059
pERK: phosphor-ERK
tERK: total ERK
TR: trained group
US: unconditioned stimulus
VEH: vehicle

## Acknowledgements

We thank Angel Vidal for the invaluable technical support. We are in debt with María Eugenia Pedreira and María Sol Fustiñana for their thorough reading of the manuscript.

## Funding

This work was supported by the following grants: ANPCYT (PICT2016 0296 and PICT2015 1199), CONICET (PIP 2014-2016 No. 11220130100519CO) and UBACYT (2018-2021 - 20020170100390BA and 2014-2017 - 20020130200283BA), Argentina.

## Notes

### Competing Interest Statement

The authors have declared no competing interest.

